# Single-cell copy number variant detection reveals the dynamics and diversity of adaptation

**DOI:** 10.1101/381590

**Authors:** Stephanie Lauer, Grace Avecilla, Pieter Spealman, Gunjan Sethia, Nathan Brandt, Sasha Levy, David Gresham

## Abstract

Copy number variants (CNVs) are a pervasive, but understudied source of genetic variation and evolutionary potential. Long-term evolution experiments in chemostats provide an ideal system for studying the molecular processes underlying CNV formation and the temporal dynamics of *de novo* CNVs. Here, we developed a fluorescent reporter to monitor gene amplifications and deletions at a specific locus with single-cell resolution. Using a CNV reporter in nitrogen-limited chemostats, we find that *GAP1* CNVs are repeatedly generated and selected during the early stages of adaptive evolution resulting in predictable dynamics of CNV selection. However, subsequent diversification of populations defines a second phase of evolutionary dynamics that cannot be predicted. Using whole genome sequencing, we identified a variety of *GAP1* CNVs that vary in size and copy number. Despite *GAP1*’s proximity to tandem repeats that facilitate intrachromosomal recombination, we find that non-allelic homologous recombination (NAHR) between flanking tandem repeats occurs infrequently. Rather, breakpoint characterization revealed that for at least 50% of *GAP1* CNVs, origin-dependent inverted-repeat amplification (ODIRA), a DNA replication mediated process, is the likely mechanism. We also find evidence that ODIRA generates *DUR3* CNVs, indicating that it may be a common mechanism of gene amplification. We combined the CNV reporter with barcode lineage tracking and found that 10^3^-10^4^ independent CNV-containing lineages initially compete within populations, which results in extreme clonal interference. Our study introduces a novel means of studying CNVs in heterogeneous cell populations and provides insight into the underlying dynamics of CNVs in evolution.

## Introduction

Copy number variants (CNVs) drive rapid adaptive evolution in diverse scenarios ranging from niche specialization to speciation and tumor evolution [1–4]. CNVs, which include gene duplications and deletions, underlie phenotypic diversity in natural populations [5–10], and can generate evolutionary novelty through modification of existing heritable material [11–14]. Beneficial CNVs are associated with defense against disease in plants, increased nutrient transport in microbes, and drug resistant phenotypes in parasites and viruses [9,15–18]. Despite the importance of CNVs for phenotypic variation, evolution and disease, the underlying dynamics with which these alleles are generated and selected in evolving populations are not well understood.

Long term experimental evolution provides an efficient means of gaining insights into evolutionary processes by allowing highly controlled and replicated selective conditions [19,20]. Chemostats are culturing devices that maintain cells in a constant nutrient-limited growth state, enabling continuous culturing [21]. Nutrient limitation in chemostats provides a defined and strong selective pressure in which CNVs have been repeatedly identified as major drivers of adaptation. CNVs containing the gene responsible for transporting the limiting nutrient are repeatedly selected in a variety of organisms and conditions including *Escherichia coli* limited for lactose [22], *Salmonella typhimurium* in different carbon source limitations [23], and *Saccharomyces cerevisiae* in glucose-, phosphate-, sulfur- and nitrogen-limited chemostats [24–30]. CNVs confer large selective advantages and multiple, independent CNV alleles have been identified within populations [25–27,31]. These findings suggest that CNVs are generated at a high rate, but estimates differ greatly, ranging from 1 x 10^−10^ to 3.4 x 10^−6^ duplications per cell per division, with variation in CNV formation rates potentially depending on locus and/or condition [32,33]. A high rate of CNV formation suggests that multiple, independent CNV-containing lineages may compete during adaptive evolution resulting in clonal interference, which is characteristic of large, evolving populations [29,34–36]. However, the extent of clonal interference among CNV-containing lineages is unknown.

The general amino acid permease, *GAP1*, is ideally suited to studying the role of CNVs in adaptive evolution. *GAP1* encodes a high-affinity transporter for all naturally occurring amino acids and analogues, and it is highly expressed in nitrogen-poor conditions [37,38]. We have previously shown that two classes of CNVs are selected at the *GAP1* locus in *S. cerevisiae*: amplification alleles in glutamine and glutamate-limited chemostats and deletion alleles in urea- and allantoin-limited chemostats [24,25]. *GAP1* CNVs are also found in natural populations. Multiple, tandem copies of *GAP1* have been identified in wild populations of the nectar yeast, *Metschnikowia reukaufii*, which result in a competitive advantage over other microbes when amino acids are scarce [39]. As a frequent target of selection in adverse environments in both experimental and natural populations, *GAP1* is a model locus for studying the dynamics and mechanisms underlying both gene amplification and deletion in evolving populations.

CNVs are generated by two primary classes of mechanisms: homologous recombination and DNA replication [40–42]. DNA double strand breaks (DSBs) are typically repaired by homologous recombination and do not result in CNV formation. However, non-allelic homologous recombination (NAHR) can generate CNVs when the incorrect repair template is used, which occurs more often with repetitive DNA sequences such as transposable elements and long terminal repeats (LTRs) [43]. During DNA replication, stalled and broken replication forks can re-initiate DNA replication through processes including break-induced replication (BIR), microhomology-mediated break-induced replication (MMBIR), and fork stalling and template switching (FoSTes) [44–46]. BIR is driven by homologous sequences, whereas MMBIR relies on shorter stretches of sequence homology. Recently, origin-dependent inverted-repeat amplification (ODIRA) has been identified as a new mechanism underlying amplification of the *SUL1* locus in yeast [47,48]. ODIRA relies on the presence of a DNA replication origin and tandem inverted repeat sequences, and is hypothesized to involve the formation of extrachromosomal circular intermediates that replicate independently.

Extrachromosomal circular DNA is common in yeast, and in humans may drive progression of tumorigenesis in cancer [49,50]. Extrachromosomal circular DNA formation is typically mediated by LTRs and other repetitive elements, and may represent an easy and reversible mechanism of generating adaptive CNVs [51,52]. Previously, we found that some *GAP1* amplifications are extrachromosomal circular elements and we hypothesized that *GAP1*^circle^ alleles are excised after NAHR between flanking LTRs [25]. Identifying the mechanisms underlying CNV formation at different loci is required for understanding the roles of CNVs in evolutionary processes and tumorigenesis.

A key limitation to the study of CNVs in evolving populations is the challenge of identifying them at low frequencies. CNVs are typically detected using molecular methods for interrogating DNA copy number including qPCR, southern blotting, DNA microarrays and sequencing [24–26]. Using any of these methods, *de novo* CNVs at a specific locus are undetectable in a heterogeneous population until present at high frequency (e.g. >50%). This precludes analysis of the early dynamics with which CNVs emerge and compete in evolving populations. As CNVs usually comprise genomic regions that include multiple neighboring genes [24], we hypothesized that CNVs could be identified on the basis of increased expression of a constitutively expressed fluorescent reporter gene inserted adjacent to a target gene of interest. A major benefit of this approach is that it detects CNVs independently of whole genome sequencing, enabling a high-resolution and inexpensive assay of CNV dynamics in evolving populations.

In this study, we constructed strains containing a fluorescent CNV reporter adjacent to *GAP1* and performed evolution experiments in different selective environments using chemostats. The CNV reporter allowed us to visualize selection of CNVs at the *GAP1* locus in real time with unprecedented temporal resolution. We find that CNV dynamics occur in two distinct phases: CNVs are selected early during adaptive evolution and quickly rise to high frequencies, but subsequent dynamics are complex. We find that *GAP1* CNVs are diverse in size and copy number, and can be generated by aneuploidy, translocations and NAHR. Nucleotide resolution of *GAP1* CNV breakpoints revealed that CNV formation is mediated by inverted repeat sequences for at least half of the resolvable cases, suggesting that replication-based mechanisms also underly gene amplification at the *GAP1* locus. Analysis of multiple *DUR3* amplifications also identified inverted repeat sequences at CNV breakpoints. These findings are consistent with ODIRA, suggesting it may be a major source of *de novo* CNVs in budding yeast. To characterize the substructure of the CNV subpopulation, we generated a lineage-tracking library using random DNA barcodes. FACS-based fractionation of CNV lineages and barcode sequencing identified thousands of individual CNV lineages consistent with a high CNV supply rate and extreme clonal interference. Together, our results show that CNVs are generated repeatedly by replication errors, and CNV-containing lineages experience extensive clonal interference, leading to complex dynamics within populations.

## Results

### Protein fluorescence increases proportionally with gene copy number

We sought to construct a reporter for CNVs that occur at a given locus of interest. Based on previous studies [53–56], we hypothesized that CNVs that alter the number of copies of a constitutively expressed fluorescent protein gene would facilitate single cell detection of *de novo* copy number variation. To test the feasibility of this approach, we constructed haploid *Saccharomyces cerevisiae* strains isogenic to the reference strain (S288c) with one or two copies of a constitutively expressed GFP variant, mCitrine [57], and diploid strains with 1-4 copies of mCitrine integrated into the genome (**Table S1**).

Flow cytometry analysis confirmed that additional copies of mCitrine produce quantitatively distinct distributions of protein fluorescence (**Figure 1A**). Haploid cells with two copies of mCitrine have higher fluorescence than those with a single copy and there is minimal overlap between the distributions of fluorescent signal in the two strains. Interestingly, normalization of the fluorescent signal by forward scatter, which is correlated with cell size, shows that the concentration of fluorescent protein is proportional to the ploidy normalized copy number of the mCitrine gene (i.e. one copy in a haploid results in a signal equivalent to two copies in a diploid and two copies in a haploid results in a signal similar to four copies in a diploid). Thus, the cell size-normalized fluorescent signal, or concentration, accurately reports on the number of copies of the fluorescent gene in single cells. Therefore, integrating a constitutively expressed fluorescent protein gene proximate to an anticipated target of selection functions as a CNV reporter for tracking gene amplifications and deletions in evolving populations (**Figure 1B**).

**Figure 1.**
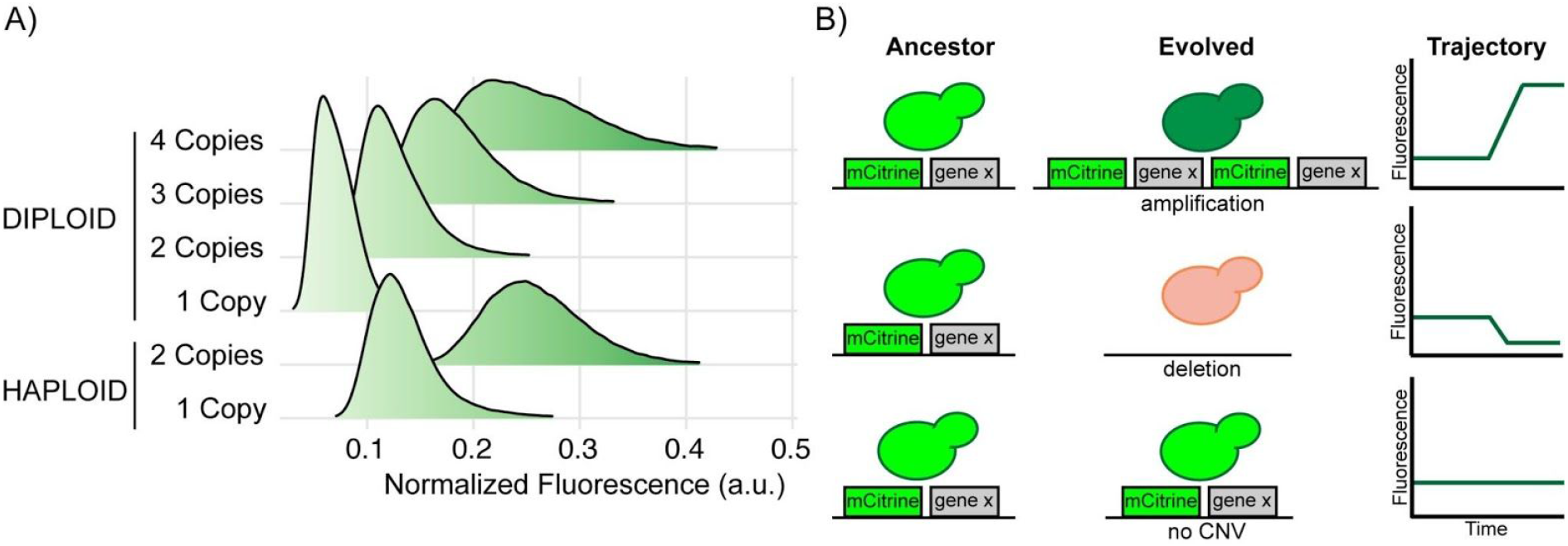
Fluorescent protein signal is proportional to gene copy number. (**A**) Protein fluorescence increases proportionally with increased copies of the mCitrine gene in both haploid and diploid cells containing variable numbers of a constitutively expressed mCitrine gene integrated at either the *HO* locus and/or the dubious ORF, *YLR123C*. The two copy diploid is heterozygous at both loci. Each distribution was estimated using 100,000 single cell measurements normalized by forward scatter, which varies as a function of cell size. (**B**) Schematic representation of how the fluorescent reporter enables CNV detection in heterogeneous evolving populations through quantitative changes in protein fluorescence.

### A CNV reporter tracks the dynamics of *GAP1* CNVs in real time

Previous work has shown that spontaneous *GAP1* amplifications are positively selected when glutamine or glutamate is the limiting nitrogen source during evolution experiments in chemostats [25]. Additionally, since *GAP1* is highly expressed in urea-limited chemostats [58] but the permease does not transport urea, *GAP1* deletions provide a fitness benefit and spontaneous deletions are selected in urea-limited conditions [25]. Thus, the use of different nitrogen sources in nitrogen-limited chemostats enables the study of both *GAP1* amplification and deletion, making it an ideal system for studying the dynamics of CNV selection in evolving populations.

We constructed a haploid strain containing a mCitrine CNV reporter 1,118 bases upstream of the *GAP1* start codon to ensure that the native regulation of *GAP1* was unaffected [59]. We inoculated the *GAP1* CNV reporter strain into 9 glutamine-, 9 urea- and 8 glucose-limited chemostats, for a total of 26 populations (**Table S2**). For each of the three selection conditions, we included two control populations: one containing a single copy of the mCitrine CNV reporter at a neutral locus (one copy control) and one containing two copies of the mCitrine CNV reporter at two neutral loci (two copy control). All populations were maintained in continuous mode (dilution rate = 0.12 culture volumes/hr; population doubling time = 5.8 hours) for 267 generations over 65 days. We sampled each of the 32 populations every 8 generations and used flow cytometry to measure fluorescence in 100,000 cells per sample.

Experimental evolution in a glutamine-limited chemostat resulted in a clear increase in fluorescence in individual cells containing the *GAP1* CNV reporter by generation 79 (**Figure 2A**). By contrast, populations containing one or two copies of mCitrine at neutral loci exhibited stable fluorescence for the duration of the experiment (**Figure 2A**). The maintenance of protein fluorescence in one and two copy control populations is consistent with the absence of a significant fitness cost associated with the CNV reporter (**Figure S1**).

**Figure 2.**
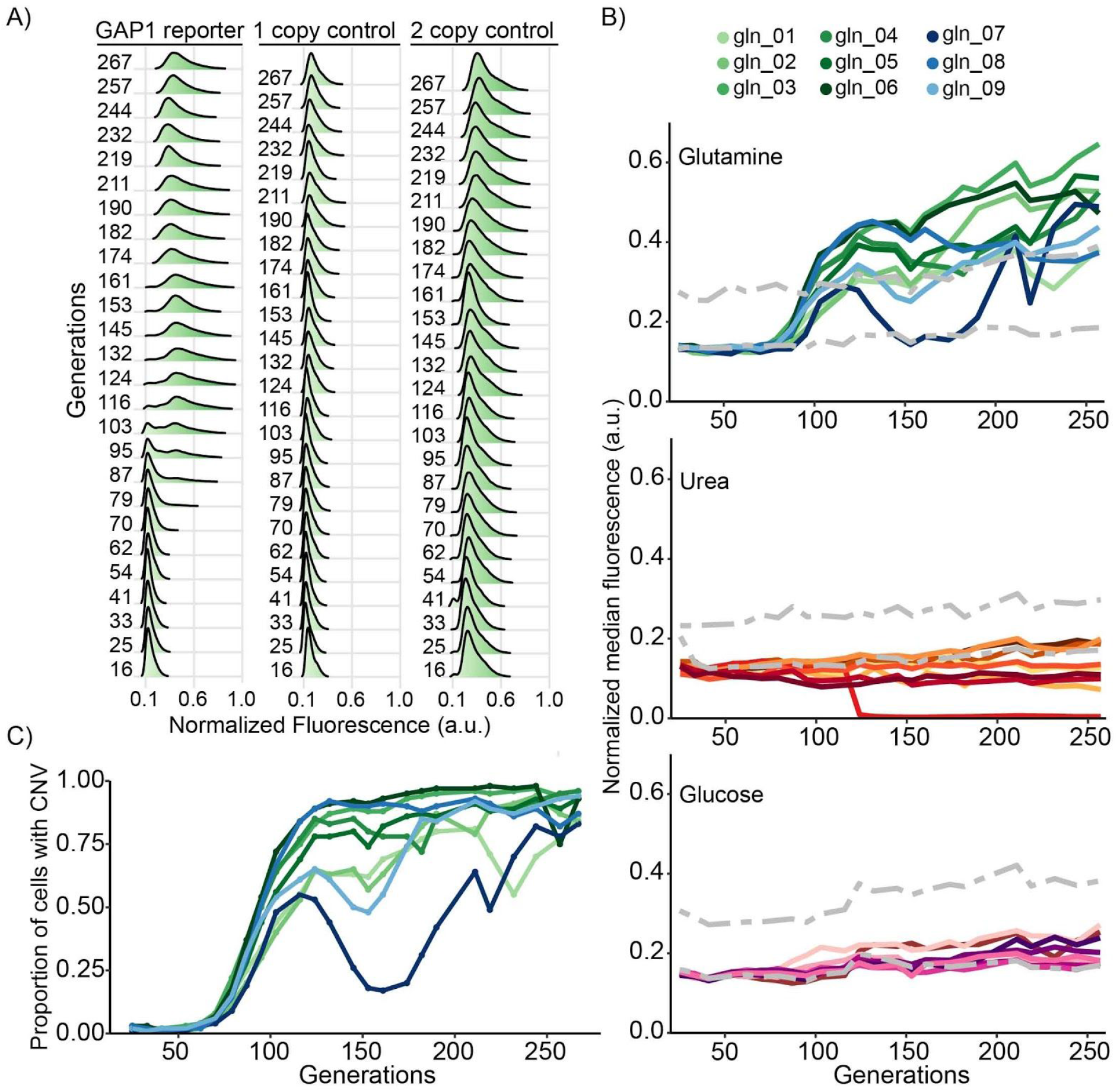
Dynamics of *GAP1* CNVs in evolving populations. (A) Normalized distributions of single-cell fluorescence over time for a representative *GAP1* CNV reporter strain and one and two copy controls evolving in glutamine-limited chemostats. Single cell fluorescence is normalized by the forward scatter measurement of the cell, which varies as a function of cell size. (B) Normalized median fluorescence for each population evolving in glutamine (n = 9), urea (n= 9) and glucose (n = 8) limited chemostats. The fluorescence of one and two copy controls are plotted for reference (grey dotted lines). (C) Estimates of the proportion of cells with *GAP1* amplifications over time for nine glutamine-limited populations containing the *GAP1* CNV reporter.

Analysis of eight additional independent populations evolving in glutamine-limited chemostats showed qualitatively similar dynamics of single-cell fluorescence over time (**Figure S2**). To summarize the dynamics of CNVs in evolving populations, we determined the median normalized fluorescence in each population at each time point. The fluorescent signal of the GAP1 CNV reporter increases with time of selection in all populations evolving in glutamine-limited chemostats (**Figure 2B**), consistent with the *de novo* generation and selection of CNVs at the *GAP1* locus in all 9 populations.

Populations evolving in urea-limited and glucose-limited chemostats do not show substantial changes in fluorescence with one exception (**Figure 2B**). In a single urea-limited population (ure_05), we detected a complete loss of fluorescent signal by generation 125, indicating the occurrence of a *GAP1* deletion that subsequently swept to fixation. Thus, the *GAP1* CNV reporter detects both amplification and deletion alleles at the *GAP1* locus. The absence of increases or decreases in fluorescence in all glucose-limited populations is consistent with the absence of selection for *GAP1* CNVs in conditions that are irrelevant for *GAP1* function.

To quantify the proportion of cells containing a *GAP1* duplication, we used one and two copy control strains to define flow cytometry gates. We found that the fluorescence of control strains varied slightly (**Figure S3A**), which may be indicative of either instrument variation or changes in cell physiology and morphology during the experiment as suggested by systematic changes in forward scatter with time (**Figure S3B**). Using a conservative method to classify individual cells containing *GAP1* amplifications (**methods**), we find that *GAP1* amplification alleles are selected with remarkably reproducible dynamics in the nine glutamine-limited populations (**Figure 2C**). CNVs are predominantly duplications (two copies), but quantification of fluorescence suggests that many cells contain three or more copies of the *GAP1* locus (**Figure S4**).

We quantified the dynamics of CNVs in each population evolved in glutamine-limited chemostats using metrics defined by Lang et al., [60]. CNVs are detected by generation 70-75 (average = 72.8 ± 2.6) in all 9 populations (T_up_) (**Table 1**). To estimate the fitness of all CNV lineages relative to the mean population fitness, we calculated S_up_, the rate of increase in the abundance of the CNV subpopulation (see **methods** and **Appendix**). The average relative fitness of the CNV subpopulation is 1.077 (S_up_) and CNV alleles are at frequencies greater than 75% in all populations by 250 generations (**Table 1**). Thus, *GAP1* amplifications emerge early, increase in frequency rapidly, and are maintained in each population for the duration of selection in glutamine-limited environments.

**Table 1.** Summary statistics of *GAP1* CNV dynamics in glutamine-limited chemostats. T_up_ is the number of elapsed generations before CNVs are reliably detected (>7% frequency, see **methods**). S_up_ is the rate of increase in CNV abundance during the initial expansion of the CNV subpopulation (**Appendix**). The frequency of CNVs in the population at generation 150 and generation 250, when genome sequencing was performed, is also indicated.

| Population | $T_{up}$ | $1 + S_{up} \pm SE$ | g150% | g250% |
| --- | --- | --- | --- | --- |
| gln_01 | 70 | $1.066 \pm 0.0038$ | 62 | 77 |
| gln_02 | 75 | $1.071 \pm 0.0034$ | 57 | 87 |
| gln_03 | 70 | $1.071 \pm 0.0037$ | 88 | 94 |
| gln_04 | 70 | $1.079 \pm 0.0036$ | 80 | 95 |
| gln_05 | 75 | $1.077 \pm 0.0041$ | 74 | 89 |
| gln_06 | 70 | $1.082 \pm 0.0043$ | 91 | 75 |
| gln_07 | 75 | $1.094 \pm 0.0048$ | 18 | 78 |
| gln_08 | 75 | $1.090 \pm 0.0052$ | 90 | 82 |
| gln_09 | 75 | $1.066 \pm 0.0050$ | 48 | 93 |
| **AVG ± STD** | **72.8 ± 2.6** | **$1.077 \pm 0.01$** | **68 ± 24** | **86 ± 8** |

We conclude that *GAP1* CNVs undergo two distinct phases of population dynamics. The initial dynamics with which CNV subpopulations emerge and increase in frequency are highly reproducible in independent evolving populations. However, after 125 generations, the trajectories of the CNV subpopulation in the different populations diverge. Many populations maintain a high frequency of *GAP1* amplification alleles, but in some populations they decrease in frequency. In one population, *GAP1* CNV alleles are nearly lost entirely from the population before subsequently increasing to an appreciable frequency (gln_07). We hypothesize that CNV lineages are selected early and reproducibly due to a high supply rate of CNVs and the strength of selection. In the second phase, the dynamics are more variable as competition between CNV lineages, other adaptive lineages, and secondary mutations influence evolutionary trajectories.

### *GAP1* CNV alleles are diverse within and between replicate populations

Based on prior studies [24,26], we hypothesized that multiple CNV alleles exist within each population. To characterize the diversity of *GAP1* CNVs, we isolated a total of 29 clones from glutamine- and 21 clones from urea-limited chemostats at 150 and 250 generations for whole genome sequencing (**Table S3**). For glutamine limitation, we enriched for clones containing *GAP1* CNVs by identifying clones with increased fluorescence. For urea limitation, we randomly selected clones. We used read depth to calculate *GAP1* copy number and to determine approximate CNV boundaries (**Figure 3A, Table S4** and **methods**). We find that *GAP1* copy number estimated by sequencing read depth correlates with the fluorescent signal for individual lineages (**Figure 3B**), indicating that fluorescent signal is predictive of copy number. In 3 clones, we find increased read depth across the entirety of chromosome XI consistent with aneuploidy. Thus, our CNV reporter is able to detect aneuploid chromosomes as well as CNVs.

**Figure 3.**
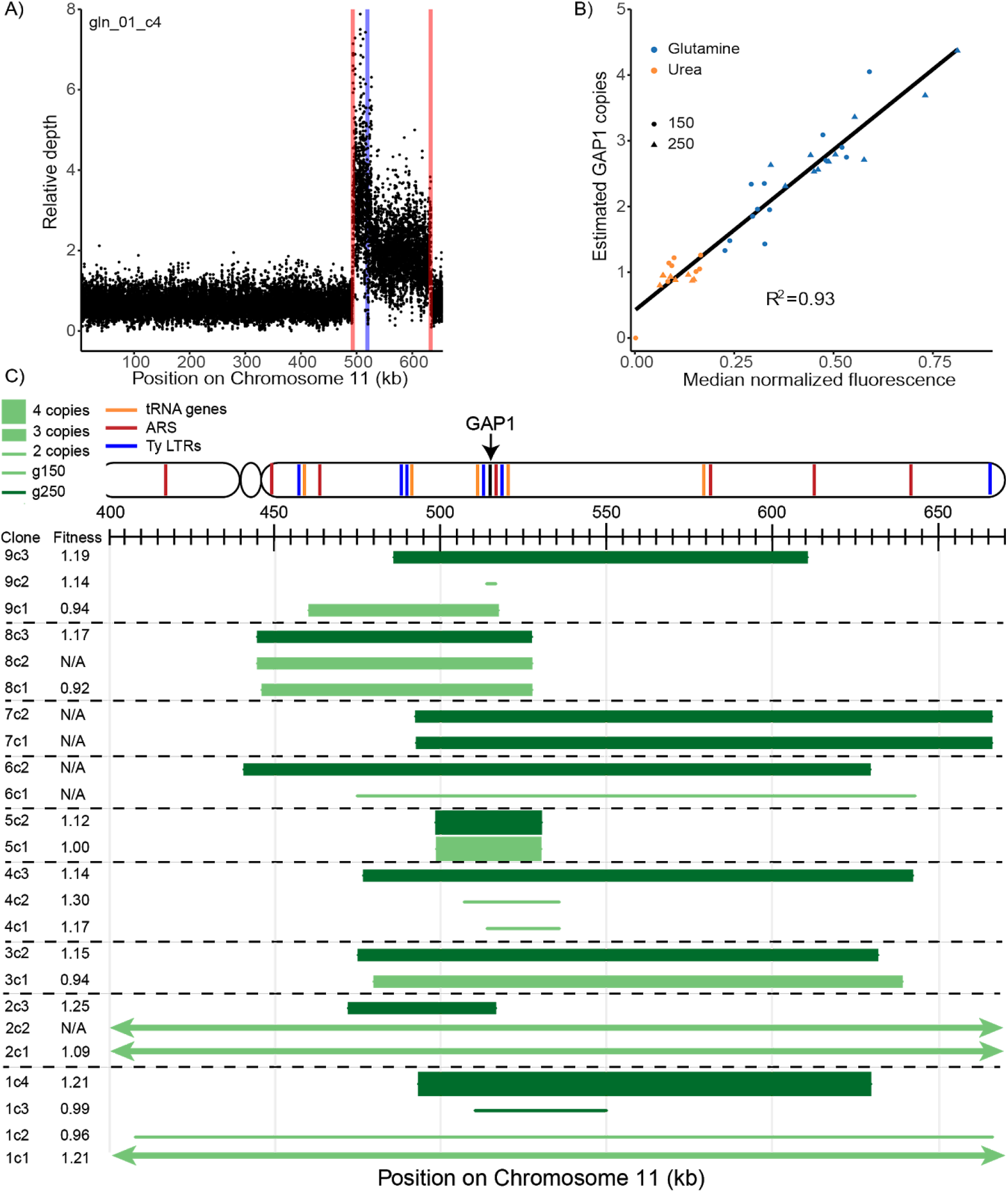
Diversity and fitness effects of *GAP1* CNVs. (A) Representative sequence read depth plot from a glutamine-limited clone (gln_01_c4). The nucleotide coordinates of *GAP1* in our CNV reporter strain are 518438-520246 (blue line). Estimated breakpoint boundaries are shown in red. Read depth was normalized to the average read depth on chromosome XI. Read depth at each nucleotide was randomly downsampled for presentation purposes. (B) Read depth based estimates of *GAP1* copy number for glutamine-limited clones are positively correlated with median fluorescence of the clonal sample, indicating that fluorescence is informative about the copy number of *de novo* CNVs. (C) Schematic representation of CNVs identified in clones isolated from glutamine-limited populations. Copy number and CNV boundaries were estimated using read depth (**methods**). The reported copy number refers specifically to the *GAP1* coding sequence and does not necessarily reflect copy number throughout the entire CNV, which may vary.

We identified diverse *GAP1* CNVs between and within populations (**Figure 3C**). Analysis of at least two *GAP1* CNV containing clones from each glumatime-limited population failed to identify a CNV common to any population. This suggests that *GAP1* CNVs are not predominantly formed through a recurrent mechanism as might be anticipated by the presence of proximate repetitive elements. Although in some cases the identical CNV was identified in different clones within populations, in the majority of populations (6/9) different clones had different CNVs. For example, at generation 150 in population gln_01 we identified a large *GAP1* CNV that includes the entire right arm of chromosome XI (gln_01_c2) and another clone that was aneuploid for the entire chromosome (gln_01). At generation 250, isolated clones from population gln_01 have CNV alleles that are distinct from each other and from those observed at generation 150. Clones from the 8 additional glutamine-limited populations show evidence for CNV diversity within and between the two time points analyzed (**Figure 3C**). Thus, sequence analysis of a small number of clonal isolates suggest the presence of multiple CNV lineages within evolving populations.

Importantly, while control populations evolving in glutamine-limited chemostats did not show evidence for *GAP1* CNVs on the basis of fluorescence, sequence analysis identified *GAP1* amplifications in lineages isolated from these populations (**Appendix**). As one and two copy control strains do not have the *GAP1* CNV reporter, this suggests that *GAP1* CNV formation and selection is not affected by the reporter. Moreover, we find no evidence that the molecular features of *GAP1* CNVs are altered as a result of the CNV reporter.

We determined the fitness of *GAP1* CNV-containing clones using pair-wise competitive fitness assays in glutamine-limited chemostats (**Figure S5** and **Figure 3C**). The majority of evolved clones (14/21) have higher relative fitness compared to the ancestral strain, with fitness coefficients ranging from 0.08-0.30, indicating that *GAP1* CNVs typically confer large fitness benefits.

### *DUR3* CNVs are repeatedly selected during urea limitation

We analyzed the genome sequences of 21 clones isolated from urea-limited populations and identified multiple CNVs at the *DUR3* locus (**Figure S6A** and **Appendix**). *DUR3* encodes a high affinity urea transporter and we have previously reported *DUR3* amplifications during experimental evolution in a urea-limited chemostat [24]. We compared properties of *GAP1* and *DUR3* amplifications and found that the average copy number for clones with *GAP1* CNVs is 3 (**Figure S6B**) whereas clones with *DUR3* CNVs contain significantly more copies with an average copy number of 5 (**Figure S6C**, t-test, p-value < 0.01). Copy number does not significantly increase in clones between 150 and 250 generations. *DUR3* CNV alleles (average of 26 kilobases) are significantly smaller than *GAP1* CNVs (average of 105 kilobases) (**Figure S6D-E**, t-test, p-value < 0.01). Thus, comparison of *GAP1* and *DUR3* CNVs suggests significant differences in the generation and selection of CNVs as a function of locus and selective condition.

### CNV breakpoints are characterized by interrupted inverted repeats

To resolve CNV breakpoint sequences, we generated a pipeline integrating CNV calls from existing CNV detection methods (CNVnator, Pindel, Lumpy, and SvABA [61–64] and optimized their performance on synthetic yeast genome data (**Supplementary Information**) simulating both clonal (**Figure S7**) and heterogeneous populations (**Figure S8**). Although these algorithms perform well using simulated data, we found that they had a high false positive and false negative rate when applied to real data (**Table S5** and **Table S6**) and in general were not informative about the novel sequence formed at CNV boundaries. Therefore, we developed a breakpoint detection pipeline that integrates information from read depth, discordant reads and split reads. To define the breakpoint sequence, we performed *de novo* assembly using all split reads and aligned the resulting contig against the reference genome (**methods**). We used information about breakpoints and copy number to infer the likely mechanism by which CNVs were generated (**methods**).

We analyzed 27 lineages containing *GAP1* CNVs and inferred the underlying mechanisms for 16 (59%) of them. Of the 16 *GAP1* CNVs that can be reliably resolved, 5 (31%) are likely the result of NAHR, 3 (19%) are the result of aneuploidies and 2 (12%) are the result of interchromosomal translocations. For 6 out of 16 (38%) *GAP1* CNVs, we identified a pair of inverted repeated sequence separated by a short distance (less than one Okazaki fragment size) proximate to the breakpoint (**Figure 4A** and **Appendix**). Typically, we find a single copy of the inverted repeat retained in the breakpoint in addition to an absent intervening sequence (AIS) that is present between the inverted repeats in the reference sequence but not the CNV allele. In some cases, breakpoints are comprised of inverted sequences that overlap in the original reference sequence. These overlaps frequently include one of the inverted repeats and are retained in the breakpoint assembly. These characteristic features comprising closely spaced inverted repeats with interstitial gain and loss, proximity to an origin of replication, and an odd copy number of amplified region are consistent with CNV formation mediated by ODIRA.

**Figure 4.**
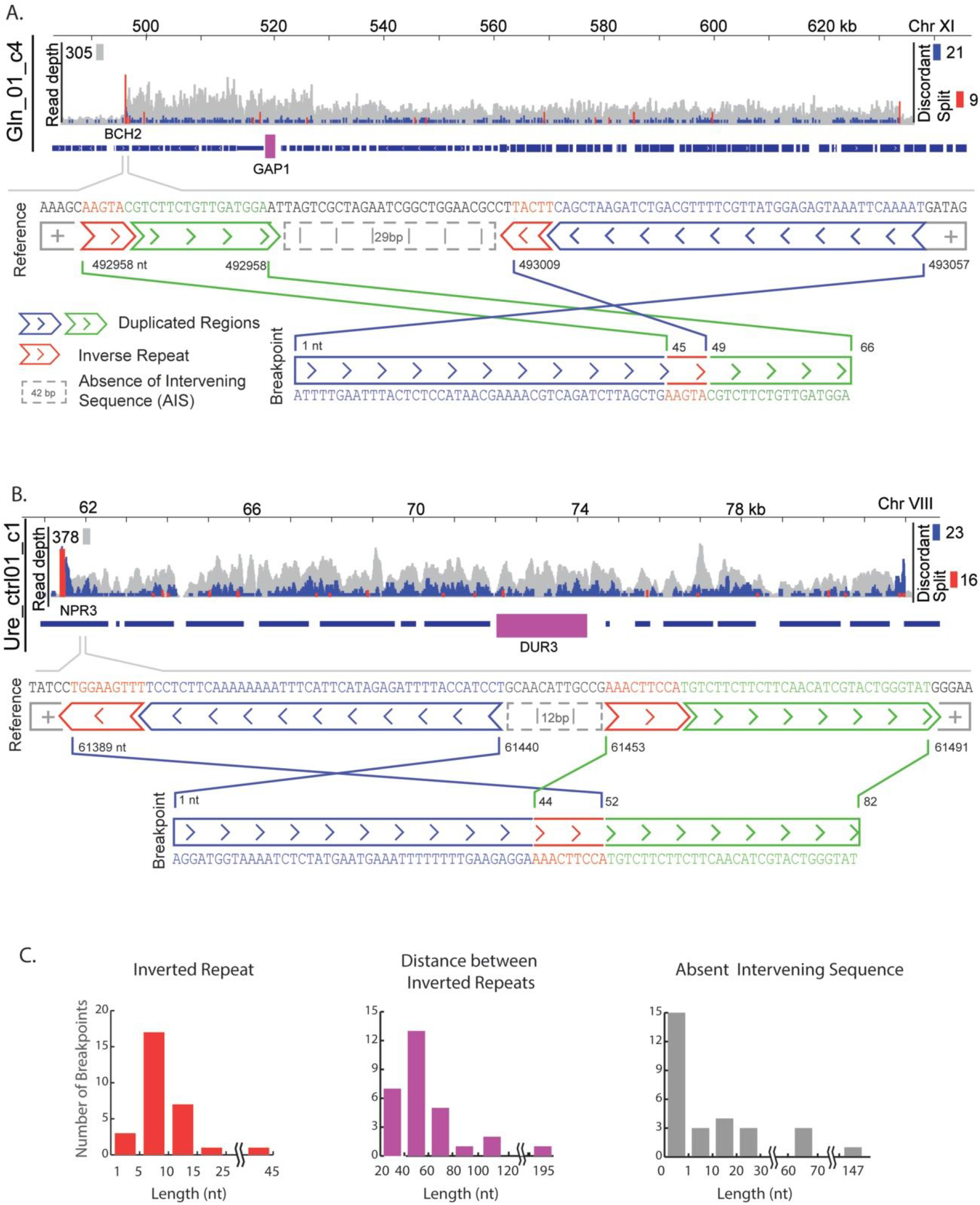
Interrupted inverted repeats mediate CNV formation. The locations of CNV breakpoints were identified using a weighted combination of co-occurrent discordant and split reads. To characterize novel sequence, we identified all supporting split reads, performed de novo assembly and aligned against the reference genome. (A) Representative CNV breakpoint for a *GAP1* CNV. The breakpoint is characterized by 5-bp inverted repeats interrupted by a 46bp sequence, which resolves to a breakpoint sequence comprising a short inverted sequence (blue) and a single copy of the inverted repeat (red). Only 17bp of the sequence between the inverted repeats is retained in the breakpoint assembly (green) with 29bp absent from the breakpoint assembly (grey). (B) Representative breakpoint of a CNV containing *DUR3*. The breakpoint sequence is characterized by a 9bp inverted repeat interrupted by a 55bp sequence. The breakpoint resolves to a 43bp inversion (blue), a single copy of the inverted repeat (red) and loss of 12 nucleotides (grey). (C) Distribution of sequence features across 29 breakpoints at the *GAP1* and *DUR3* loci that contain inverted repeats.

Analysis of *DUR3* amplifications suggests that all *DUR3* CNVs are formed through ODIRA (**Table S5**). *DUR3* amplification breakpoints occur at inverted repeat sequences and typically result in an AIS (**Figure 4B** and **Appendix**). Analysis of all ODIRA-mediated breakpoints shows that the inverted repeat sequences range in length from 4-15bp with a median length of 8bp that are spaced 50bp apart (**Figure 4C**). In those cases that result in an AIS at the breakpoint, the absent sequence is typically 20bp (**Figure 4C**).

### Whole genome population sequencing provides insight into population heterogeneity

To comprehensively characterize acquired genomic variation in populations, we performed whole population, whole genome sequencing of glutamine-, urea-, and glucose-limited populations at generations 150 and 250 (**Table S3**). Analysis of relative sequence read depth is consistent with high frequency *GAP1* CNVs in glutamine-limited populations (see **Appendix**). Population sequencing also confirmed the fixation of a *GAP1* deletion (ure_05) in a urea-limited population. Relative sequence read depth at the *GAP1* locus correlates well with the normalized fluorescence of the *GAP1* CNV reporter in populations (**Figure S9**) providing additional evidence for the utility of the CNV reporter. In glutamine-limited chemostats, *GAP1* copy number estimated within populations (which is a function of copy number within clones and allele frequencies) ranges from 2-4 copies, with a trend towards increased copy number over time (**Figure S9**).

We performed single nucleotide polymorphism (SNP) analysis using genome sequencing data from populations (**Table S7**) and clones (**Table S8**) at generations 150 and 250. More non-synonymous SNPs were identified in glucose-limited populations than glutamine- and urea-limited populations (**Table 2**), which contained *GAP1* and *DUR3* amplifications at high frequencies at 150 and 250 generations. In contrast to previous studies [28,29], we did not identify CNVs at the *HXT6/7* locus in glucose-limited populations. Therefore, increased nucleotide variation within these populations may reflect alternative adaptive strategies in glucose-limited populations.

**Table 2.** Summary of single nucleotide variation in three different selection conditions. Populations were sequenced at 150 and 250 generations. For variants that were identified at both time points, we determined whether they increased (↑) or decreased (↓) in frequency between generation 150 and 250.

|  | Glucose (n=10) |  |  |  | Urea (n=11) |  |  |  | Glutamine (n=11) |  |  |  |
| --- | --- | --- | --- | --- | --- | --- | --- | --- | --- | --- | --- | --- |
|  | Total |  | Trend |  | Total |  | Trend |  | Total |  | Trend |  |
| Predicted effect | g150 | g250 | ↑ | ↓ | g150 | g250 | ↑ | ↓ | g150 | g250 | ↑ | ↓ |
| Non-coding | 4 | 9 | 2 | 1 | 8 | 12 | 1 | 0 | 6 | 5 | 0 | 0 |
| Missense | 47 | 61 | 17 | 4 | 22 | 34 | 6 | 2 | 12 | 22 | 4 | 1 |
| Frameshift | 4 | 5 | 1 | 1 | 5 | 6 | 2 | 0 | 2 | 2 | 0 | 0 |
| Synonymous | 1 | 7 | 0 | 0 | 4 | 10 | 2 | 0 | 4 | 3 | 0 | 0 |
| Stop gained | 4 | 10 | 2 | 0 | 7 | 4 | 0 | 2 | 0 | 5 | 0 | 0 |
| Start lost | 0 | 1 | 0 | 0 | 0 | 0 | 0 | 0 | 0 | 0 | 0 | 0 |
| Splice variant | 2 | 3 | 0 | 1 | 0 | 0 | 0 | 0 | 0 | 1 | 0 | 0 |
| Mitochondrial genome | 0 | 0 | 0 | 0 | 0 | 2 | 0 | 0 | 0 | 0 | 0 | 0 |
| Inframe insertion | 0 | 0 | 0 | 0 | 0 | 0 | 0 | 0 | 0 | 1 | 0 | 0 |
| Total variants | 62 | 96 | 22 | 7 | 46 | 68 | 11 | 4 | 24 | 39 | 4 | 1 |

We find multiple genes with independent, non-synonymous variation in glutamine-limited populations (**Table 3**) including *MCK1*, a protein kinase with potential roles in NHEJ; *SOG2*, a member of the RAM signaling pathway and regulator of bud separation after mitosis; and *TAO3*, another member of the RAM network. Our lab has previously reported mutations in *MCK1* during glutamine and arginine limitation [24], suggesting that it is a recurrent target of selection in these conditions. Changes in cell morphology are potentially adaptive in nutrient-poor conditions, which may result from defects in cell cycle progression and bud separation associated with mutations in the RAM pathway [65].

**Table 3.** Genes with multiple, independent, non-synonymous acquired mutations. Variants with frequency greater than 5% within each population.

| Glucose-limitation |  | Urea-limitation |  | Glutamine-limitation |  |
| --- | --- | --- | --- | --- | --- |
| Gene Name | Total Variants | Gene Name | Total Variants | Gene Name | Total Variants |
| *TRK1* | 11 | *DUR1,2* | 14 | *MCK1* | 3 |
| *SVF1* | 2 |  |  | *SOG2* | 3 |
| *CDC48* | 3 |  |  | *TAO3* | 2 |
| *WHI2* | 3 |  |  | *GPB2* | 2 |

In the nine urea-limited populations, we identified 14 independent non-synonymous variants in *DUR1,2* (**Table 3**). *DUR1,2*, encodes urea amidolyase, which metabolizes urea to ammonium. At two different locations in the gene, we find that the exact nucleotide was mutated multiple times independently. In a third location, we identified a SNP at the exact nucleotide position as previously reported [24]. Thus, a subset of variants in *DUR1,2* appear to be uniquely beneficial and recurrently selected. Interestingly, we did not detect nonsense mutations among these 14 mutations suggesting that acquired variants alter *DUR1,2* function rather than rendering it non-functional.

In glucose-limited populations, we identified multiple, independent mutations in four genes (**Table 3**): *TRK1*, a component of the potassium transport system; *SVF1*, which is important for the diauxic growth shift and is implicated in cell survival during aneuploidy [66]; *CDC48*, an AAA ATPase; and *WHI2*, which is a mediator of the cellular stress response. Previous studies have identified loss-of-function mutations in *WHI2* suggesting it is a general target of selection across different conditions [24,27,67].

Analysis of clonal samples (**Table S8**) was largely consistent with population sequencing. We identified two cases in which SNPs occurred within *GAP1* CNVs. These SNPs are present at frequencies of 53% in a lineage containing a *GAP1* duplication and 30% in a lineage containing a *GAP1* triplication indicating that they are present on only one of the copies within the CNV. We also identified polymorphisms within *DUR3* amplifications. This suggests that individual copies of a gene within a CNV can accumulate additional mutations even in relatively short-term evolutionary scenarios. Interestingly, 8 of the 9 clones with *DUR3* amplifications also acquired a variant in *DUR1,2*, which may be indicative of a synergistic relationship between CNVs and single nucleotide variation.

### Lineage tracking reveals extensive clonal interference among CNV lineages

The reproducible dynamics of CNV lineages observed during glutamine-limited experimental evolution may be due to two non-exclusive reasons: 1) a high supply rate of *de novo* CNVs or 2) pre-existing CNVs in the ancestral population (**Figure S10**). In both scenarios, a single CNV or multiple, competing CNVs may underlie the reproducible dynamics. Sequence analysis of clonal lineages suggests at least two, and as many as four, CNV lineages may co-exist in populations (**Figure 3**); however, genome sequencing is uninformative about the total number of lineages for two key reasons. First, the potential for recurrence of CNVs confounds distinguishing CNVs that are identical by state from those that are identical by descent. Second, as gene amplifications can exhibit instability, CNVs that arise *de novo* may subsequently diversify over time resulting in apparently distinct alleles that are derived from a common event.

To quantify the number, relationship and dynamics of individual CNV lineages, we constructed a lineage tracking library using random DNA barcodes [68]. We constructed a library of ~80,000 unique barcodes (**Figure S11**) in the background of the *GAP1* CNV reporter and performed six independent replicates in glutamine-limited chemostats. Real time monitoring of CNV dynamics using the *GAP1* CNV reporter recapitulated the dynamics of our original experiment (**Figure 5A, Figure S12A** and **Table S9**) although CNV lineages appeared earlier in these populations (T_up_; t-test p-value < 0.01). As the lineage tracking strain was independently derived from the strain used in our original experiment, these results indicate that selection of *GAP1* CNVs in glutamine limitation is reproducible and independent of strain background.

**Figure 5.**
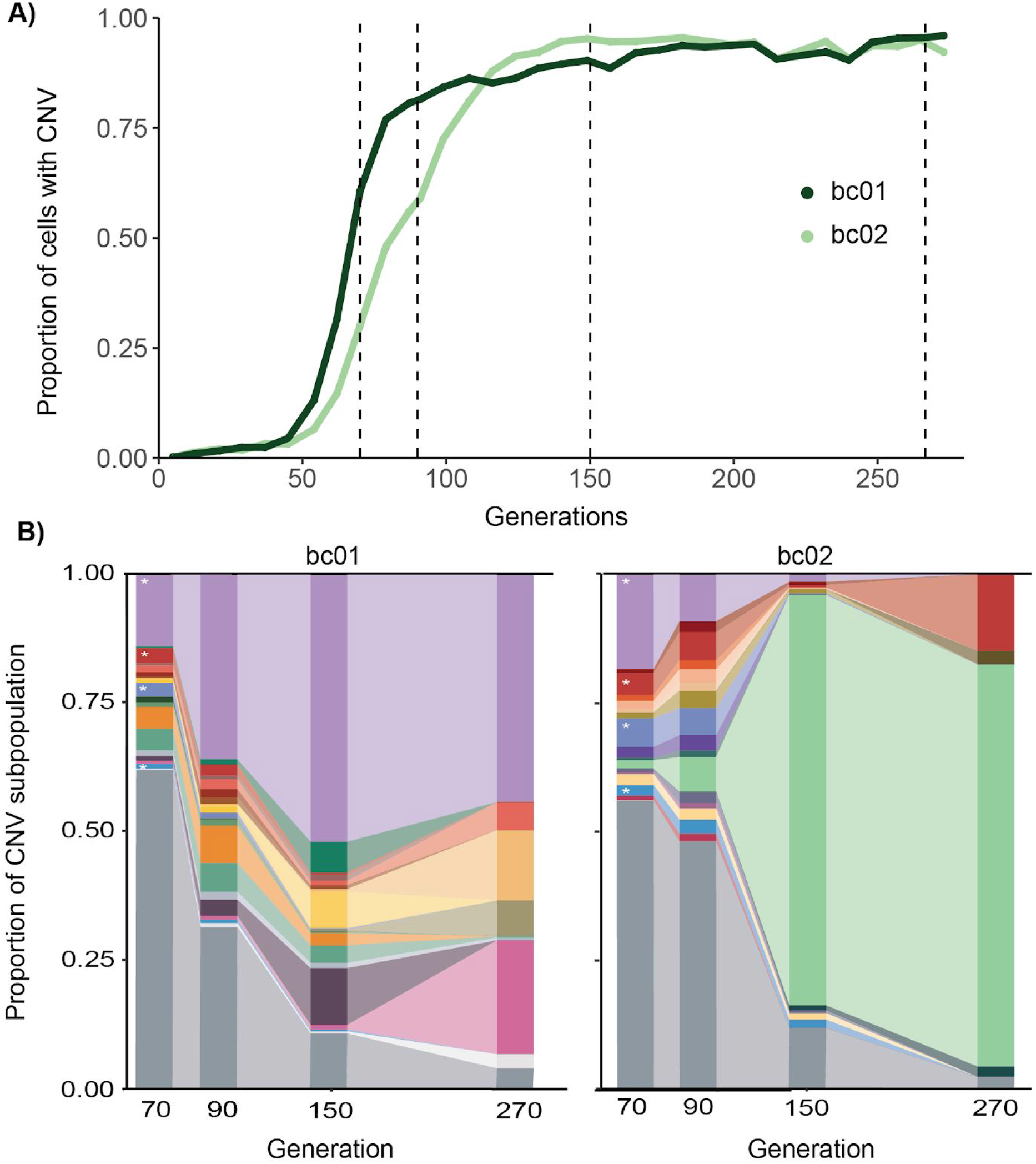
Lineage tracking reveals extensive clonal interference among CNV-containing lineages. (**A**) We used fluorescence-activated cell sorting (FACS) to fractionate cells containing *GAP1* CNVs from 2 independent populations at four time points (dashed black lines) and performed barcode sequencing. (**B**) Using a sample- and time point-specific false positive correction, we identified 7067, 973, 131, and 76 barcodes in bc01 (right) at generations 70, 90,150 and 270 respectively and 5305, 5351, 583, and 28 barcodes in bc02 (left) at generations 70, 90,150 and 270 respectively. The 21 unique barcodes in bc01 and 18 unique barcodes in bc02 that are present at >1% population frequency in at least one time point are represented by a unique color in the plot. All other lineages that are never found at >1% frequency are shown in grey. Lineages denoted by a * are found at >1% frequency in both populations.

To quantify individual lineages, we isolated the subpopulation containing CNVs from two populations (bc01 and bc02) at multiple timepoints (generations 70, 90, 150, and 270) using fluorescence activated cell sorting (FACS) (**Figure 5A**). We sequenced barcodes from the CNV subpopulation at each time point and determined the number of unique lineages ([69] and **methods**). To account for variation in the purity of the isolated CNV subpopulation, we analyzed individual clones from the CNV subpopulation isolated by FACS to estimate a false positive rate, which we find varies as a function of time point (**Figure S12B** and **methods**), and applied this correction to barcode counts (**Table S10**).

Strikingly, we detect thousands of independent *GAP1* CNV lineages at generation 70 indicating that a large number of independent *GAP1* CNVs are generated and selected in the early stages of glutamine limitation (**Figure 5B**). Applying our false positive correction, we identified 7,067 *GAP1* CNV lineages in bc01 and 5,305 GAP1 CNV lineages in bc02 at generation 70 (**methods; Table S10**). To increase the stringency of CNV lineage identification, we required a lineage tracking barcode be detected in at least two consecutive time points in the CNV subpopulation. Using this conservative criterion, we detected 872 and 2,695 CNV-containing lineages in bc01 and bc02, respectively (**Table S10**). Thus, between 10^3^-10^4^ independent CNV lineages initially compete within each population containing ~10^8^ cells.

The overall diversity of CNV lineages decreases dramatically with time, consistent with decreases in lineage diversity observed in other evolution experiments [68,70]. By generation 270, we detect only 76 CNV lineages in bc01 and 28 CNV lineages in bc02. To determine the dominant lineages, we identified barcodes that reached greater than 1% frequency in the CNV subpopulation in at least one time point: 21 independent lineages are found at greater than 1% frequency in bc01 and 18 independent lineages are found at greater than 1% frequency in bc02 (**Figure 5B**). These results indicate the presence and persistence of multiple *GAP1* CNVs, where glutamine-limited populations are characterized by large initial diversity followed by a dramatic reduction in the diversity of CNV lineages.

Although CNVs rise to high population frequencies in both populations (**Figure 5A**), the composition of competing CNV lineages is dramatically different: in bc02, a single lineage dominates the population by generation 150 (**Figure 5B**), whereas in bc01, there is much greater diversity at these latter time points. In both populations, several CNV lineages that comprise a large fraction of the CNV subpopulation at early generations (generations 70, 90, or 150) are extinct by generation 270. Thus, within populations, individual CNV lineages do not increase in frequency with uniform dynamics despite consistent and reproducible dynamics of the entire CNV subpopulations (**Figures 5A** and **Figure 2**). Differences in fitness between individual CNV lineages, possibly as a result of copy number, CNV size and secondary adaptive mutations, are likely to contribute to these dynamics.

### CNV subpopulations comprise *de novo* and pre-existing CNV alleles

To distinguish the contribution of pre-existing genetic variation and *de novo* variation to CNV lineage dynamics, we assessed whether barcodes were shared between CNV lineages in independent populations. We identified four barcodes at greater than 1% frequency that are common to both populations (**Figure 5B**) and are therefore likely to have been present in the ancestral population. One of these barcodes (indicated in light purple) was present at 14% in bc01 and 19% in bc02 at generation 70. We find that the barcode for this lineage was over-represented in the ancestral unselected population (an initial frequency of 0.014%, which is one order of magnitude greater than the average starting frequency of 0.0011%; **Figure S11**). Although there is a possibility that *de novo* CNVs formed independently in this barcode lineage, it is more likely that this lineage contained a pre-existing CNV in the ancestral population. While this lineage represented a sizable fraction of the CNV subpopulation in both replicate populations, it was only maintained at high frequency in bc01. Only one of the four pre-existing CNV lineages persists throughout the experiment in both populations. By contrast, in each population, we identified 17 and 14 unique high frequency CNV lineages that are most likely new CNVs. These results indicate that both pre-existing CNVs and *de novo* CNVs that arise during glutamine limitation contribute to adaptive evolution.

## Discussion

Copy number variants are a primary class of genetic variation and adaptive potential. In this study, we sought to understand the short-term fate of CNVs as they are generated and selected in evolving populations. Previous work from our laboratory and others has shown that well-defined, strong selection in chemostats provides an ideal system for studying CNVs. We used nitrogen limitation to establish conditions that positively select for amplification and deletion of the gene *GAP1*, which encodes the general amino acid permease, in *S. cerevisiae*.

### A *GAP1* CNV reporter reveals the dynamics of selection

To determine the dynamics with which CNVs are selected at the *GAP1* locus, we used a constitutively expressed fluorescent gene located adjacent to *GAP1* and tracked changes in single cell fluorescence over time. While one and two copy control strains with *mCitrine* at neutral loci maintain a steady fluorescent signal over 250 generations of selection, all glutamine-limited populations with the *GAP1* CNV reporter show increased fluorescence by generation 75. Importantly, the structure and breakpoints of CNVs within and between populations are different, indicating independent formation of CNVs. Control strains were inoculated independently, and have different genetic backgrounds, but also form CNVs at the *GAP1* locus as determined by whole genome sequencing. These data indicate that *GAP1* CNVs are positively selected early and repeatedly in glutamine-limited environments.

### Inferences of CNV formation mechanisms

Whole genome sequencing of *GAP1* CNV lineages that were isolated on the basis of increased fluorescence uncovered a wide range of CNVs within and between populations. We found cases in which distinct alleles were identified within populations at different time points and cases in which we identified the same CNV allele 100 generations later. *GAP1* CNV alleles are 105 kilobases on average, but can include the entire right arm of chromosome XI (260 kilobases). A previous study in bacteria showed that there is a cost to gene duplication, with a fitness reduction of 0.15% per kilobase [71]. Therefore, we hypothesized that CNVs would decrease in size over evolutionary time through a refinement process in order to reduce the fitness burden. However, we failed to detect a significant reduction in CNV allele size over time. This may be because increased CNV size does not confer a fitness cost in yeast, the fitness benefit of the *GAP1* CNV outweighs this cost, or because there are other genes within the CNV whose amplification confers a fitness benefit.

Breakpoint analysis provided insight into CNV formation mechanisms. We identified breakpoints within LTRs for 5 glutamine-limited clones. These clones all have 2 copies of *GAP1*, which suggests that these CNVs were formed by a tandem duplication mediated through NAHR. Surprisingly, *GAP1* gene amplifications occurring between the flanking LTRs *YKRCdelta11* and *YKRCdelta12* are infrequent (3/27 *GAP1* CNVs). The *GAP1* deletion, which occurred in one population in urea limitation, had breakpoints in *YKRCdelta11* and *YKRCdelta12*. We did not find evidence for the selection of *GAP1^circle^* CNVs in any population.

We identified 6 *GAP1* CNVs and 9 *DUR3* CNVs containing a pair of breakpoints that comprise closely-spaced inverted repeat sequences. These CNVs have specific characteristics that are consistent with the ODIRA mechanism, including an odd copy number and proximity to an autonomously replicating sequence. Thus, while NAHR is likely to play a role in CNV formation, our results suggest that replication-based mechanisms may be a predominant source of gene amplifications in yeast.

The large number of CNV lineages identified in our study indicates that they occur at a high rate. Recent studies have suggested that adaptive mutations are stimulated by the environment. Stress can lead to increases in genome-wide mutation rates in both bacteria and yeast [72–74] and replicative stress can lead directly to increased formation of CNVs [75,76]. Other groups have proposed an interplay between transcription and CNV generation, and that active transcription units might even be “hotspots” [76]. These hotspots, often designated as common fragile sites, may occur in large, late replicating genes, and be a function of inter-origin distance [77–79]. Local transcription at the rDNA locus leads to rDNA amplification, and is thought to be regulated in response to the environment [80]. Transcription of the *CUP1* locus in response to environmental copper leads to promoter activity that further destabilizes stalled replication forks and generates CNVs [81]. Given the high level of *GAP1* transcription in nitrogen limited chemostats [58] it is tempting to speculate that this condition may promote the formation of *GAP1* CNVs. Further studies are required to understand the full extent of processes that underlie CNV formation at the *GAP1* locus and how these different mechanisms may contribute to the fitness and overall success of CNV lineages.

### Clonal interference underlies CNV dynamics

By combining a CNV reporter with lineage tracking, we identified a surprisingly large number of independent CNV lineages. Whereas clonal isolation and sequencing suggested at least four independent lineages, lineage tracking indicates that many thousands of individual CNV lineages emerge within less than 100 generations. Most of these lineages do not ever achieve high frequencies in the populations, as we identified only 18-21 lineages present at >1% frequency. We hypothesize that the frequency of *GAP1* CNV events can be attributed to a combination of factors including: a high mutation supply rate due in part to the large chemostat population size (~10^8^), the strength of selection, and the fitness benefit conferred by *GAP1* amplification. Together, these factors contribute to an early, deterministic phase, during which CNVs are formed at a high rate and thousands of lineages with CNVs rapidly increase in frequency. During a second phase, the dynamics are more variable as competition from different types of adaptive lineages, and additional acquired variation, influence evolutionary trajectories of individual CNV lineages. This phenomenon has recently been observed in other evolution experiments, where early events are driven by multiple competing single-mutant lineages [70], but later dynamics are influenced by stochastic factors and secondary mutations [68].

While we found lineages that were common to both populations (at least one of which is likely to contain a pre-existing CNV), ancestral lineages do not exclusively drive the dynamics we observe. Pre-existing CNV lineages have different dynamics in each population, and do not prevent the emergence of unique *de novo* CNV lineages. This demonstrates that the ultimate fate of a CNV lineage depends on multiple factors, and a high frequency at an early generation does not guarantee that a lineage will persist in the population. Thus, CNV dynamics result from pre-existing and *de novo* variation and are characterized by extensive clonal interference and replacement among competing CNV lineages.

### Conclusion

The combined use of a fluorescent CNV reporter and barcode lineage tracking provides unprecedented insight into this important and understudied class of mutations. Previous studies have tracked specific mutations and their fitness effects [60], but ours is the first single-cell based approach to identify an entire class of mutations and follow their evolutionary trajectories with high resolution. While barcode tracking alone would provide information about the number of adaptive lineages present and their fitness effects, the CNV reporter enables us to explicitly determine the number of unique CNV events. Rather than identifying CNVs that are identical by descent, the barcodes enable a quantitative estimate of clonal interference. In addition, the reporter provides an estimate of the total proportion of CNVs in the population, which we can use to inform our understanding of lineage dynamics. Our system provides a facile means for studying the molecular processes underlying CNV generation as well as evolutionary aspects of CNVs including pleiotropy and tradeoffs. Extension of our method is likely to be useful for addressing fundamental questions regarding the evolutionary and pathogenic role of CNVs in diverse systems.

## Methods

### Strains and media

We used FY4 and FY4/5, haploid and diploid derivatives of the reference strain S288c, for all experiments. **Table S1** is a comprehensive list of strains constructed and used in this study. To generate fluorescent strains, we performed high efficiency yeast transformation [82] with an mCitrine gene under control of the constitutively expressed *ACT1* promoter (*ACT1pr::mCitrine::ADH1term*) and marked by the KanMX G418-resistance cassette (*TEFpr::KanMX::TEFterm*). The entire construct, which we refer to as the mCitrine CNV reporter, is 3,375 base pairs. For control strains, the mCitrine reporter was integrated at two neutral loci: *HO* (*YDL227C*) on chromosome IV and the dubious ORF, *YLR123C* on chromosome XII. Diploid control strains containing 3 and 4 copies of the mCitrine CNV reporter were generated using a combination of backcrossing and mating. We constructed the *GAP1* CNV reporter by integrating the mCitrine construct at an intergenic region 1 kilobase upstream of *GAP1* (integration coordinates, chromosome XI: 513945-517320). PCR and Sanger sequencing were used to confirm integration of the *GAP1* CNV reporter at each location (PCR primer sequences are provided in **Table S11**). Transformants were backcrossed, sporulated, and segregants genotyped to identify the correct genotype.

For the purpose of lineage tracking, we constructed a strain containing a landing pad and the *GAP1* CNV reporter by mating and segregation analysis [68]. As the kanMX cassette is present at two loci in this cross, we performed tetrad dissection and identified four spore tetrads that exhibited 2:2 G418 resistance. A segregant with the correct genotype (G418 resistant, ura-) was identified and confirmed using a combination of PCR and fluorescence analysis. We introduced a library of random barcodes by transformation and selection on SC-ura plates [68]. We plated an average of 500 transformants on 200 petri plates and estimated 78,000 independent transformants.

Nitrogen limiting media (glutamine and urea limitations) contained 800 μM nitrogen regardless of molecular form and 1 g/L CaCl_2_-2H_2_O, 1 g/L of NaCl, 5 g/L of MgSO_4_-7H_2_O, 10 g/L KH_2_PO_4_, 2% glucose and trace metals and vitamins as previously described [24]. Glucose limiting media contained 0.08% glucose, 1 g/L CaCl_2_-2H_2_O, 1 g/L of NaCl, 5 g/L of MgSO_4_-7H_2_O, 10 g/L KH_2_PO_4_, 50g/L (NH_4_)_2_SO_4_ and trace metals and vitamins [83].

### Long-term experimental evolution

We inoculated the *GAP1* CNV reporter strain into 20mL ministat vessels [84] containing either glutamine-, urea-, or glucose-limited media. Control populations containing either one or two copies of the CNV reporter at neutral loci (*HO* and *YLR123C*) were also inoculated in ministat vessels for each media condition. Ministats were maintained at 30°C in aerobic conditions and diluted at a rate of 0.12 hr^−1^ (corresponding to a population doubling time of 5.8 hours). Steady state populations of 3 x 10^8^ cells were maintained in continuous mode for 270 generations (~2 months). Every 30 generations, we archived 2 mL population samples at −80°C in 15% glycerol.

### Flow cytometry sampling and analysis

To monitor the dynamics of CNVs, we sampled 1mL from each population every ~8 generations. We performed sonication to disrupt any cellular aggregates and immediately analyzed the samples on an Accuri flow cytometer, measuring 100,000 cells per population for mCitrine fluorescence signal (excitation = 516nm, emission = 529nm, filter = 514/20nm), cell size (forward scatter) and cell complexity (side scatter). We generated a modified version of our laboratory flow cytometry pipeline for this analysis (https://qithub.com/GreshamLab/flow), which uses the R package *flowCore* [85]. We used forward scatter height (FSC-H) and forward scatter area (FSC-A) to filter out doublets, and FSC-A and side scatter area (SSC-A) to filter debris. We quantified fluorescence for each cell and divided this value by the forward scatter measurement for the cell to account for differences in cell size. To determine population frequencies of cells with zero, one, two, and three plus copies of *GAP1*, we used one and two copy control strains grown in the same media to define gates and perform manual gating. We used a conservative gating approach to reduce the number of false positive CNV calls by first manually drawing a liberal gate for the one copy control strain, followed by a non-overlapping gate for the two copy control strain.

### Quantification of CNV dynamics

To quantify the dynamics of CNVs in evolving populations, we defined summary statistics as in [60]. T_up_ is the generation at which CNVs are initially detected and S_up_ is the slope of the linear fit during initial population expansion of CNVs. We first determined the proportion of cells with a CNV and the proportion of cells without CNVs at each time point, using the manually defined gates. To calculate T_up_, we defined a false positive rate for CNV detection in evolving one copy control strains from generations 1-153 (defined as the average plus one standard deviation = 7.1%). We designate T_up_ once an experimental population surpasses this threshold. To calculate S_up_, we plotted the natural log of the ratio of the proportion of cells with and without a CNV against time and calculated the linear fit during initial population expansion of CNVs. We defined the linear phase on the basis of R^2^ values (**Appendix**). S_up_ can also be defined as the percent increase in CNVs per generation, which is an approximation for the relative average fitness of all CNV alleles in the population.

### Isolation and analysis of evolved clones

Clonal isolates were obtained from each glutamine- and urea-limited population at generation 150 and generation 250. We isolated clones by plating cells onto rich media (YPD) and randomly selecting individual colonies. We inoculated each clone into 96 well plates containing the limited media used for evolution experiments and analyzed them on an Accuri flow cytometer following 24 hours of growth. We compared fluorescence to unevolved ancestral strains and evolved 1 and 2 copy controls grown under the same conditions, and chose a subset of clones for whole genome sequencing (**Table S4**).

To measure the fitness coefficient of evolved clones, we performed competitive fitness assays in glutamine-limited chemostats using the same conditions as our evolution experiments [24]. We co-cultured our fluorescent evolved strains with a non-fluorescent, unevolved reference strain (FY4). We determined the relative abundance of each strain every 2-3 generations for approximately 15 generations using flow cytometry. We performed linear analysis of the natural log of the ratio of the two genotypes against time and estimated the fitness, and associated error, relative to the ancestral strain.

### Genome sequencing

For both population and clonal samples, we performed genomic DNA extraction using a modified Hoffman-Wnston protocol [86]. We used SYBR Green I to measure gDNA concentration, standardized each sample to 2.5 ng/μL, and constructed libraries using tagmentation following a modified Illumina Nextera library preparation protocol [87]. To perform PCR clean-up and size selection, we used an Agilent Bravo liquid handling robot. We measured the concentration of purified libraries using SYBR Green I and pooled libraries by balancing their concentrations. We measured fragment size with an Agilent TapeStation 2200 and performed qPCR to determine the final library concentration.

DNA libraries were sequenced using a paired-end (2×75) protocol on an Illumina NextSeq 500. Standard metrics were used to assess data quality (Q30 and %PF). To remove reads from a potentially contaminating organism that was introduced after recovery from the chemostats, we filtered any reads that aligned to *Pichia kudriavzevii*. Given the evolutionary divergence between these species, the majority of filtered reads belonged to rDNA and similar, deeply conserved sequences. The median percent contamination was 1.165%. We modified the S. cerevisiae reference genome from NCBI (assembly R64) to include the entire *GAP1* CNV reporter and aligned all reads to this reference. We aligned reads using bwa mem ([88], version 0.7.15) and generated BAM files using samtools ([89], version 1.3.1). Summary statistics for all sequenced samples are provided in **Table S3**. FASTQ files for all sequencing are available from the SRA (accession SRP142330).

### CNV detection using published algorithms

To assess the performance of CNV detection algorithms, we simulated copy number variants ranging in size from 50bp - 100,000bp in 100 synthetic yeast genomes. We used SURVIVOR [90] to simulate copy number variants in the reference yeast genome and wgsim [89] to generate corresponding paired-end FASTQ files. We used bwa mem [88] to map reads back to the reference and called CNVs with Pindel, CNVnator, LUMPY and SvABA [61–64]. We assessed the effect of read depth on algorithm performance by downsampling a 100x coverage BAM file to 80x, 50x, 20x, 10x and 5x coverage. We defined a CNV as being correctly predicted if the simulated and detected CNVs were: (i) of the same type (e.g. duplication), (ii) predicted to be on the same chromosome and (iii) contained in the same interval (defined by the start and stop position), which were considered overlapping if there was no gap between them (maxgap=0) and had minimum overlap of one base pair (minoverlap=1). For intervals [a,b] and [c,d], where a<=b and c<=d, when c<=b and d>=a, the two intervals overlap and when c > b or d < a, the two intervals don’t overlap. If the gap between these two intervals is <= maxgap and the length of overlap between these two intervals is >= minoverlap, the two intervals are considered to be overlapping.

To assess the performance of these tools on heterogeneous population samples, we also simulated mixed samples by combining reads from a simulated CNV-containing genome and an unmodified reference yeast genome at varying proportions. The ratio of the reads from the CNV-containing genome varied between 20% and 90%, and the total coverage was 50X.

Performance comparisons for all benchmarking were based on false discovery rate (FDR) and F-score. The F-score (also known as F1 measure) combines sensitivity/recall(r) and precision(p) with an equal weight using the formula F=(2pr)/(p+r). An F-score reaches its best value at 1 and worst at 0 and was multiplied by 100 to convert to a percentage value. We called CNVs for each clone and population sample using an in-house pipeline that collates results from Pindel, SvABA, and Lumpy (**Table S5** and **Table S6**).

### Sequence read depth and breakpoint analysis

To manually estimate CNVs boundaries we used a read-depth based approach. For each sample sequenced, we used samtools [89] to determine the read depth for each nucleotide in the genome. We liberally defined CNVs by identifying ≥300 base pairs of contiguous sequence when read depth was ≥3 times the standard deviation across chromosome XI for *GAP1* or chromosome VIII for *DUR3*. These boundaries were further refined by visual inspection of contiguous sequence ≥100 base pairs with read depth ≥3 times the standard deviation. These analyses were only performed on sequenced clones because population samples are likely to have multiple CNVs and breakpoints thereby confounding read-depth based approaches. We compared manually estimated breakpoints to those identified by the algorithms (**Table S5**) and defined a set of ‘high confidence breakpoints’.

To determine CNV breakpoints, we extracted split and discordant reads from bam files using samblaster [91]. Both split reads and discordant reads were used to identify breakpoints; however, each candidate breakpoint required at least one split read. Additionally, breakpoints were scored using a weighted method where split reads were worth 1 and discordant reads were worth 3. Positively identified breakpoints required a score of at least 9. Breakpoint sequences were generated by making local assemblies of breakpoint associated split and discordant reads using velvet [92]. The relationship between breakpoint sequences and the reference genome was determined using BLAST.

To infer the underlying mechanism by which CNVs were formed, we applied the following criteria. If the sequence of least one of the two CNV boundaries contained inverted repeat sequences, and we estimated an odd number of copies in the CNV, we classified the mechanism as ODIRA. If the boundary sequences occurred within repetitive sequence elements (LTRs or telomeres) and had two copies we inferred NAHR. Aneuploids were defined on the basis of increased read depth throughout the entire chromosome, but no detected novel sequence junctions. Translocations were identified by LUMPY. All breakpoints that failed to meet these criteria were defined as unresolved.

### SNP and variant identification

SNPs and indel variants were first identified using GATK4’s Mutect2 [93], which allows for the identification of variants in evolved samples (‘Tumor’) after filtering using matched unevolved samples (‘Normal) and pool of normals (PON). The PON was constructed using six sequenced ancestral clones, while the paired normal was a single, deeply sequenced, ancestor. Variants were further filtered using GATK’s FilterMutectCalls to remove flag low quality predictions; only variants flagged as passed or germline risk were retained. We chose to retain so-called germline risk variants because this flag is often raised when a variant is at or near 100% frequency in the evolved sample but can not definitively be ruled out from being present in the ancestral population. Given the haploid nature of the evolved population, and further downstream filtering of ‘too-recurrent’ mutations, we allowed germline risk variants to be retained. Variants were further filtered if they occurred in low complexity sequence; i.e. variants were filtered if the SNP or indel occurred in, or generated, a homogenous nucleotide stretch of five or more of the same nucleotide. Variants from within populations that were detected at less than 5% frequency were considered low confidence and excluded. Finally, variants were filtered if they were found to be ‘too-recurrent’; that is, if the exact nucleotide variant was identified in more than three independently evolved lineages, we deemed it more parsimonious to assume that the variant was present in the ancestor.

### Quantifying the number of CNV lineages

We inoculated the lineage tracking library into 20mL ministat vessels [84] containing glutamine-limited media. Control populations containing either zero (WT), one or two copies of the *GAP1* CNV reporter at neutral loci (HO and YLR123C) were also inoculated in ministat vessels for each media condition. Control populations did not contain lineage tracking barcodes. Ministat vessels were maintained and archived as above. Samples were taken for flow cytometry every ~8 generations and analyzed as previously described.

We used fluorescence activated cell sorting (FACS) to isolate the subpopulation of cells containing two or more copies of the the mCitrine CNV reporter using a FACSAria. We defined our gates using zero, one, and two copy mCitrine control strains sampled from ministat vessels at the corresponding timepoints: 70, 90, 150, and 265 generations. Depending on the sample, we isolated 500,000-1,000,000 cells with increased fluorescence, corresponding to two or more copies of the reporter. We grew the isolated subpopulation containing CNVs for 48 hours in glutamine-limited media and performed genomic DNA extraction using a modified Hoffman-Winston protocol [86]. We verified FACS isolation of true CNVs by isolating clones from sub-populations sorted at generation 70, 90, and 150 (sorted from all lineage tracking populations, bc01-06) and performing independent flow cytometry analysis using an Accuri. We estimated the average false positive rate of CNV isolation at each time point as the percent of clones from a population with FL1 less than one standard deviation above the median FL1 in the one copy control strain. Only subpopulations with fluorescence measurements for at least 25 clones were included in calculations of false positive rate.

We performed a sequential PCR protocol to amplify DNA barcodes and purified the products using a Nucleospin PCR clean-up kit [68]. We quantified DNA concentrations by qPCR before balancing and pooling libraries. DNA libraries were sequenced using a paired-end (2x×150) protocol on an Illumina MiSeq 300 Cycle v2. Standard metrics were used to assess data quality (Q30 and %PF, **Table S3**). However, the reverse read failed due to over-clustering, so all analyses were performed only using the forward read. We used the Bartender algorithm with UMI handling to account for PCR duplicates and to cluster sequences with merging decisions based solely on distance except in cases of low coverage (<500 reads/barcode), for which the default cluster merging threshold was used [69]. Clusters with a size less than four or with high entropy (>0.75 quality score) were discarded. We estimated relative abundance of barcodes using the number of unique reads supporting a cluster compared to total library size.

## Supplementary Note

### Application of existing CNV detection algorithms fails to identify *GAP1* and *DUR3* duplications

The identification of CNVs using a fluorescent reporter enables assessment of the extent to which current algorithms are able to detect CNVs using short read sequencing. To evaluate algorithm performance, we first tested each algorithm using simulated data (**methods**). We simulated haploid clonal samples and found that LUMPY, SvABA and Pindel performed reasonably well (FDR < 5% and F-score > 80%) with average genome coverages ranging from 5X-50X (**Figure S7**). We then simulated mixed non-clonal populations containing CNVs at differing allele frequencies. We found that LUMPY and SvABA performed well in all scenarios whereas Pindel performed well only when simulated populations contained CNVs at frequencies of at least 50% (**Figure S8**).

On the basis of these results, we generated an in-house pipeline for detecting CNVs that integrates results from Pindel, Lumpy, and SvABA [61–63]. We first applied this pipeline to clonal CNV samples with increased fluorescence. We found that in 36% of cases, in which a CNV is predicted to be present on the basis of fluorescence and clear increases in sequencing read depth, all three algorithms failed to call a CNV at the GAP1 locus. LUMPY consistently identified the most breakpoints, which we compared against manual CNV boundary calls (**Table S5**). This high false negative rate was further exacerbated for population sequencing data in which fluorescence clearly indicated the presence of CNV lineages at >50% frequencies: in 28% of tested populations a CNV was not called by any of the algorithms (**Table S6**). We conclude that existing CNV calling algorithms are poorly suited to detection of CNVs in heterogeneous samples and suffer from a high false negative rate even for clonal haploid samples.

## Supplementary Figures

**Figure S1. Assessment of CNV reporter fitness effects**. The fitness of strains carrying one (DGY500) or two copies (DGY1315) of a constitutively expressed mCitrine gene was assayed. Fluorescent strains were co-cultured with the non-fluorescent, unevolved reference strain (FY4). We performed two independent competitive fitness assays in glutamine-limited chemostats using the same conditions as evolution experiments. No significant fitness defect was observed for either strain indicating that constitutive expression of the fluorescent gene does not confer a fitness cost in these conditions. Error bars are 95% confidence intervals.

**Figure S2. The *GAP1* CNV reporter indicates the emergence of *GAP1* CNVs in all glutamine-limited populations**. Distributions of single-cell fluorescence over time for all glutamine-limited experimental populations. Fluorescent signal is normalized by forward scatter, which varies as a function of cell size. Each distribution is based on at least 100,000 single cell measurements.

**Figure S3. Normalization by forward scatter mitigates effects of cell physiology and morphology variation on CNV reporter signal**. Dashed grey lines represent one and two copy control populations. (**A**) Median unnormalized fluorescence across time for all evolving populations. (**B**) Median forward scatter over time for all populations. One glucose-limited population (pink) developed a bud separation defect, resulting in a cell aggregation phenotype and large forward scatter and fluorescence measurements. Normalizing by forward scatter accounts for this issue and other changes in overall cell physiology during the evolution experiments (see **Figure 2B**).

**Figure S4. Gating flow cytometry data enables estimation of CNV alleles that contain more than two copies**. The proportion of cells with zero, one, two, and three or more copies of *GAP1* in each glutamine-limited experimental population. This proportion was calculated after generating gating criteria based on one and two copy control populations.

**Figure S5. *GAP1* CNV-containing lineages have a higher relative fitness than the ancestral strain**. The fitness of evolved lineages containing *GAP1* CNVs was determined by competition experiments with a non-fluorescent, unevolved reference strain (FY4) in glutamine-limited chemostats. The majority (14/21) of evolved CNV-containing lineages have significantly higher fitness (t-test, p.val < 0.05) than the ancestor. Decreased, or insignificant fitness differences, may reflect context-specific fitness effects of *GAP1* CNV-containing lineages. Error bars are 95% confidence intervals.

**Figure S6. Identification of CNV alleles at the *DUR3* locus.** (**A**) A schematic illustrating the genomic context and estimated breakpoints for clones isolated from urea-limited chemostats containing *DUR3* CNVs. Breakpoint boundaries were estimated using a read depth based approach (**methods**). Compared to (**B**) clones isolated from glutamine-limited chemostats containing *GAP1* CNVs, (**C**) clones isolated from urea-limited chemostats have significantly higher copy number. (**D**) *GAP1* CNV alleles are significantly larger than (**E**) *DUR3* CNV alleles (t-test p.val < 0.01).

**Figure S7. Benchmarking existing CNV detection algorithms with simulated clonal samples**. We simulated CNVs in the yeast genome at different average sequencing depths to assess the performance of CNVnator, LUMPY, Pindel and SvABA. Algorithm performance was evaluated using false discovery rate (FDR) and F-score. We find that with increased read depth (**A**) the false discovery rate increases for deletion detection, but (**B**) overall performance improves for all algorithms as determined by F-score. Conversely, for duplication detection (**C**) the false positive rate is not increased with increasing read depth and (**D**) overall performance improves with increased read depth.

**Figure S8. Benchmarking existing CNV detection algorithms with simulated heterogeneous populations samples**. We simulated heterogeneous populations containing CNVs at varying frequencies and assessed algorithm performance. Most algorithms perform reasonably well when CNVs are present at 50% or higher in the population.

**Figure S9. Population estimates of *GAP1* copy number by CNV reporter and quantitative sequencing are linearly correlated and increase with time of adaptive evolution**. Relative depth at the *GAP1* locus, calculated from whole genome sequencing data, is strongly correlated with the median normalized fluorescence of the *GAP1* CNV reporter in populations. Glutamine-limited populations measured at generation 250 tend to have higher fluorescence and higher relative read depth at the *GAP1* locus than at generation 150.

**Figure S10. Population prehistory of independent evolution experiments**. All independent populations share a common history prior to founding of individual populations. The prehistory of experiments using the GAP1 CNV reporter (**A**) differ with respect to the size of the size founding population in experiments using a lineage tracking library (**B**). Any variation that is introduced prior to founding of individual populations may contribute to the evolution of all populations. Variation that is introduced after separation into individual populations contributes to evolutionary outcomes in that population only.

**Figure S11. Distribution of barcodes counts in ancestral populations**. We determined the distribution of read counts supporting each unique barcode in the ancestral population, after filtering out low confidence clusters. The relative frequencies of barcodes vary by over an order of magnitude and we observe a long tail with a few barcodes significantly overrepresented in the ancestral population. The red arrow indicates an overrepresented barcode in the ancestral population that was identified in the CNV subpopulation in both independent barcoded evolution experiments (indicated in purple in **Figure 5B**).

**Figure S12. Identification of barcoded *GAP1* CNV-lineages in evolving populations**. (**A**) *GAP1* CNV dynamics in barcoded populations assayed using a CNV reporter. (**B**) Estimation of true positive rate of CNV isolation by FACS at generations 70, 90, and 150. CNV subpopulations were isolated by FACS at each timepoint and clones isolated by plating for single colonies. The percent of cells containing a CNV in the fractionated subpopulation was estimated using at least 25 clones. A one copy control strain was used to define gates.

## Supplementary Tables

**Table S1**. List of strains used and generated in this study.

**Table S2**. List of all experimentally evolved populations.

**Table S3**. DNA sequencing summary statistics for all clonal and population samples.

**Table S4**. Summary statistics of all evolved clones. Nucleotide resolution of estimated break points and size (in kilobases) of CNV alleles are also presented. These statistics were calculated using a read-depth based approach. RD = read depth.

**Table S5**. Breakpoint analysis of 27 *GAP1* CNVs and 9 *DUR3* CNVs. We compare all 3 CNV detection methods used in this study: breakpoint sequences determined through split read assembly and alignment, breakpoint identification using Lumpy, and CNV boundary classification using using read-depth and visual inspection.

**Table S6**. Summary of CNV detection algorithm performance for all population samples.

**Table S7**. SNPs identified from population sequencing data. Mode refers to whether a mutation detected at both times points and increased in frequency (increase) or decreased in frequency (decrease). SNPs present at frequencies greater than 0.05 are reported.

**Table S8**. SNPs identified from clone sequencing data. SNPs were filtered on the basis of their frequency in the clonal sequence data using a threshold of 0.25.

**Table S9**. Summary statistics for *GAP1* CNV dynamics, determined using the *GAP1* CNV reporter, in replicated evolution experiments using lineage tracking libraries. Summary statistics are defined as in Table 1.

**Table S10**. Frequency of barcodes in each population across time. We estimated the number of GAP1 CNV containing lineages by correcting the number of barcodes identified by the estimated false positive rate of CNV isolation by FACS. High confidence *GAP1* CNV lineages are defined as those that are found at two or more timepoints.

**Table S11**. List of primers used to generate PCR products for strain construction and to confirm insertion of PCR products after transformation.

## Acknowledgements

We thank members of the Gresham, Vogel and Hochwagen labs for helpful discussions. This work was supported by NIH grant R01GM107466 and NSF grant MCB1244219 to DG and a NSF GRFP DGE1342536 to GA.

